# Biogeographical patterns in soil bacterial communities across the Arctic region

**DOI:** 10.1101/655431

**Authors:** Lucie A Malard, Muhammad Zohaib Anwar, Carsten S Jacobsen, David A Pearce

**Affiliations:** Faculty of Health and Life Sciences, Northumbria University, Newcastle-upon-Tyne NE1 8ST, United Kingdom; Department of Environmental Sciences, Aarhus University, 4000 Roskilde, Denmark; British Antarctic Survey, High Cross Madingley Road, Cambridge, CB3 0ET, United Kingdom

**Keywords:** 16S rRNA, Arctic soil, microbial diversity, indicator species, core microbiome, biogeography

## Abstract

The considerable microbial diversity of soils, their variety and key role in biogeochemical cycling has led to growing interest in their global distribution and the impact that environmental change might have at the regional level. In the broadest study of Arctic soil bacterial communities to date, we used high-throughput DNA sequencing to investigate the bacterial diversity from 200 independent Arctic soil samples from 43 sites. We quantified the impact of spatial and environmental factors on bacterial community structure using variation partitioning analysis, illustrating a non-random distribution across the region. pH was confirmed as the key environmental driver structuring Arctic soil bacterial communities, while total organic carbon, moisture and conductivity were shown to have little effect. Specialist taxa were more abundant in acidic and alkaline soils while generalist taxa were more abundant in acidoneutral soils. Of 48,147 bacterial taxa, a core microbiome composed of only 13 taxa that were ubiquitously distributed and present within 95% of samples was identified, illustrating the high potential for endemism in the region. Overall, our results demonstrate the importance of spatial and edaphic factors on the structure of Arctic soil bacterial communities.

## Introduction

Frozen soils in the Arctic region store over 1500 Pg of carbon (Koven et al., 2011; Mackelprang et al., 2011) and as Arctic warming is exacerbated and permafrost thaw accelerates, the depth of the active layer is increasing. As previously frozen carbon becomes available, it is expected that microbial activity will increase, which may lead to increased atmospheric release rates of climate active gases such as carbon dioxide (CO_2_), methane (CH_4_) and nitrous oxide (N_2_O) (Ma et al., 2007; Mackelprang et al., 2011). Carbon-climate feedback studies of permafrost affected regions use temperature, soil moisture and precipitation as the main drivers of decomposition rates (Koven et al., 2011; Schuur et al., 2015). While models are useful to gain a global understanding of the impact of climate change on permafrost thaw and greenhouse gas release, the accuracy of results obtained is highly variable when compared with data collected in the field or laboratory (Schuur et al., 2015) due to empirical and modelling uncertainties which still need to be addressed (Bradford et al., 2016).

As microorganisms drive biogeochemical cycling and participate in the uptake and release of CO_2_, CH_4_ and N_2_O, microbial data should be incorporated in climate models. Current models use soil properties to model changes in fluxes, without considering microbial communities and the changes in community composition induced by climate change (Bardgett et al., 2008; Nazaries et al., 2013). Adding microbial information into models will improve their predictions; however, detailed microbial data is still required, with a focus on the microbial community, diversity, function, stability and long-term changes in these communities (Graham et al., 2012; Nazaries et al., 2013). While global surveys of microbial diversity have already been conducted (Tedersoo et al., 2014; Delgado-Baquerizo et al., 2018), the number of Arctic samples is restricted (Malard and Pearce, 2018) and therefore, microbial data is still lacking for most permafrost-affected regions.

To accurately model spatial distribution patterns of microbes and incorporate microbial data from permafrost-affected regions in climate models in the near future, the first step is to produce a baseline database of microbial diversity with respect to biogeographical distribution. Biogeography is the study of biodiversity across space and time, it gives insights into ecological mechanisms such as speciation, extinction, dispersal and species interactions (Martiny et al., 2006; Fierer, 2008), improving our understanding of community assembly. Theoretically, distant and isolated habitats are expected to present high endemicity as a consequence of intrinsic dispersal limitations and environmental filtering (Mittelbach and Schemske, 2015; Kleinteich et al., 2017; Bahram et al., 2018). Thus, isolated, pristine ecosystems with limited human presence, such as the Arctic region, should harbour endemic communities. However, microbial communities may be less constrained by geographical barriers and thus, have long been considered ubiquitous (Finlay, 2002; O’Malley, 2007). Yet, recent studies have uncovered patterns of microbial biogeography on global scales (Fierer and Jackson, 2006; Lauber et al., 2009; Tedersoo et al., 2014; Henschel et al., 2015; Bahram et al., 2018; Delgado-Baquerizo et al., 2018). They demonstrated that microbial distribution is non-random, generally influenced by biotic and abiotic factors such as plant cover, soil pH, C/N ratio or precipitation, and that communities vary from one region to another. However, whether these patterns apply to Arctic microbial communities is still subject to debate as these studies tend to have a low number of Arctic samples and Arctic studies have generally focused on small scales patterns in restricted areas, as shown by Metcalfe et al. (2018) highlighting the focuses on Abisko, Sweden and Toolik lake, Alaska. Large scale studies are therefore required to improve our understanding not only of the spatial distribution of microorganisms but also on the processes of community assembly. Producing a baseline database of diversity in the region will enable the accurate modelling of spatial distribution, linking with patterns of functional processes to evaluate the potential consequences of environmental change on ecosystem properties.

In this study, we conducted a Pan-Arctic survey of bacterial communities in Arctic soils to provide a baseline of Arctic bacterial diversity. We addressed the following questions: (i) Are bacterial taxa ubiquitously distributed or are there biogeographical patterns ? (ii) What are the key factors influencing bacterial community structure ? (iii) Are there taxa closely associated with the key edaphic properties and what can we learn from them? (iv) If bacterial taxa are not ubiquitous, is there a core microbiome?

## Methods

### Sample collection

Soil samples were collected across the Arctic region between April 2017 and September 2017, the GPS coordinates of each site were recorded with a portable GPS [Fig. 1]. All sites were generally remote with limited human presence; however, in Iceland and Norway where tourist activities are frequent, we aimed to sample at least 500 m away from human-related activities. At each location, 3 to 5 soil samples were collected within 100 m^2^ under the most common vegetation, for a total of 200 independent Arctic samples across 43 sites, with the aim to cover the broadest geographical area possible. Despite the extent, it should be noted that large areas of Canada and Russia remain uncharacterized and under-represented, highlighting areas where sampling should be prioritized in the future. Approximately 150 g of soil per sample was collected from the top 15 cm, avoiding plant roots and rocks and using an ethanol-cleaned shovel and Whirl-Pak bags (Nasco, WI, USA). Remaining plant roots and rocks were removed manually in a class II microbiological safety cabinet; samples were homogenized by manual mixing and frozen at −20 °C before transportation to the United Kingdom. Samples were conserved at – 20 °C until analysed.

**Figure 1:**
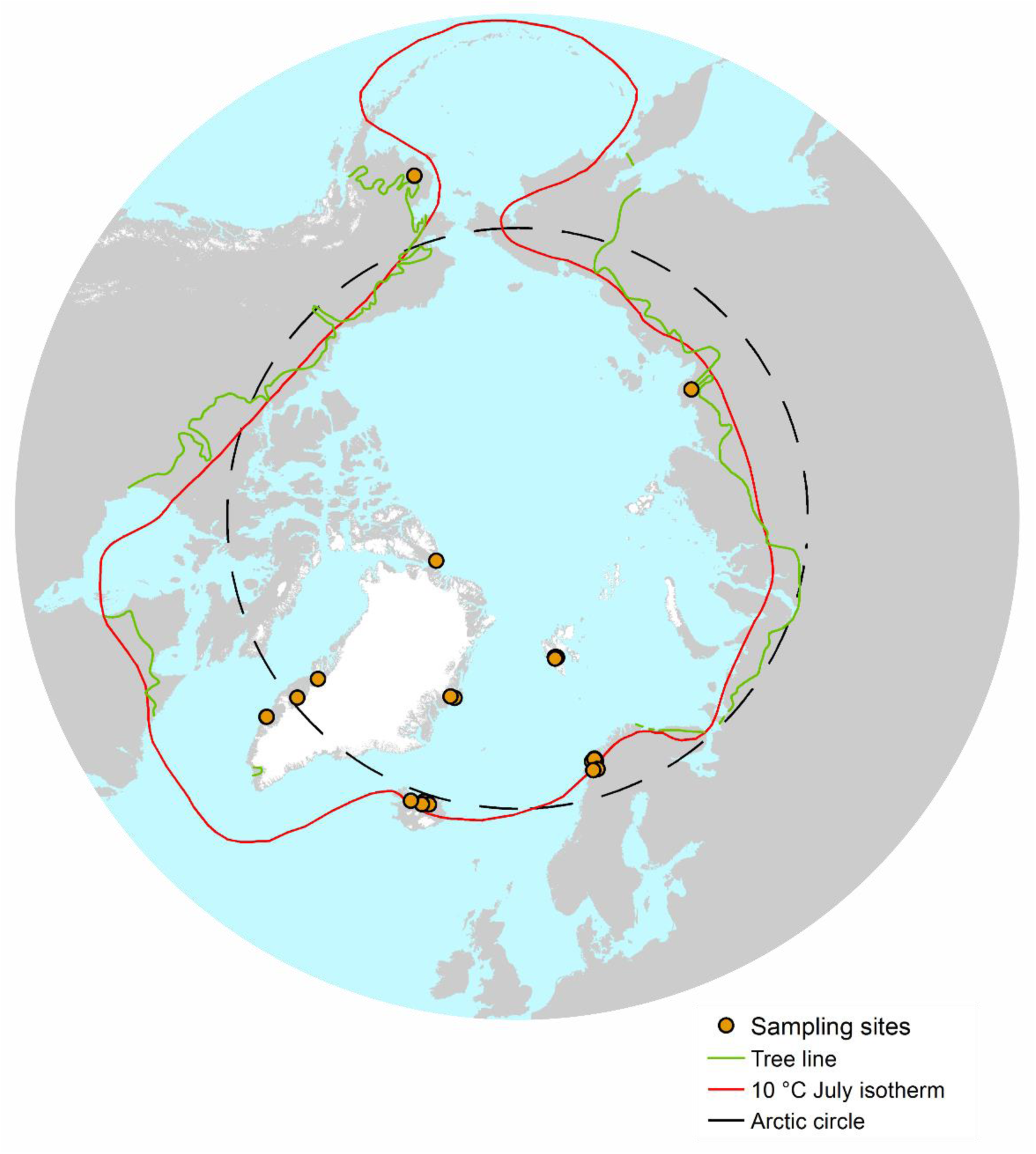
Map of sampling sites within the Arctic Region, as characterised by the Arctic circle (black line), the 10 °C isotherm (red) and the tree line boundary (green). 200 individual soil samples at 43 different sites were collected for this study, all within at least one of the definition of the Arctic region.

### Soil properties

We focused our investigation on environmental variables previously identified as influencing bacterial communities and included pH, conductivity, moisture and organic carbon. Moisture content was measured gravimetrically for each soil sample after drying at 150 °C for 24 h and total organic content (TOC) was measured gravimetrically by heating previously dried soils to 550 °C for 4 h. pH and conductivity were measured in the laboratory in a 1:5 freshly thawed soil to water ratio, using a Mettler-Toledo FE20 pH meter (Mettler-Toledo Instruments co., Shanghai, China) and a CMD500 conductivity meter (WPA, Cambridge, UK).

### DNA extraction

Soil DNA was extracted for each sample using the PowerSoil kit (Qiagen, Hilden, Germany), for a total of 400 DNA extracts. Each sample was PCR amplified using the universal primers 515F-806R, as per the Schloss lab standard operating Procedure (Kozich et al., 2013) and the Earth Microbiome Project (Thompson et al., 2017), under the following conditions: initial denaturation at 95°C for 2 min then 30 cycles of 20 s denaturation at 95°C; primer annealing at 55°C for 15 s; elongation at 72°C for 5 min then a final elongation at 72°C for 10 min. Negative controls, DNA extraction kit controls and ZymoBIOMICS mock communities (Zymo Research, Irvine, CA, USA) were included alongside the soil DNA and sequenced. PCR amplicons were cleaned and normalized using SequalPrep Plate Normalization Kit (Invitrogen, Carlsbad, CA, USA) and combined into four pools. Each pool was quantified using fragment size determined by BioAnalyzer hsDNA assay (Agilent technologies, Santa Clara, CA, USA) and concentration by Qubit hsDNA kit (Invitrogen). The library was supplemented with 5% PhiX and loaded on an Illumina MiSeq V2 500 cycles cartridge.

### Illumina Sequencing and Data Processing

Raw amplicon sequences were demultiplexed with the associated barcodes. Cutadapt (Martin, 2011) was used for adaptor and primer clipping. Forward and reverse reads longer than 240 bp were merged (98 % ± 0.8 % / sample) using FLASH (fast length adjustment of short reads) (Magoč and Salzberg, 2011) for a total of 20 million reads (∼50,000 ± 30,000 reads/sample) initially. Vsearch (Rognes et al., 2016) was used for downstream analyses. Quality filtering was carried out with an expected error < 1.5. Dereplication was performed to identify unique sequences. A two-step chimera detection method was used, first by aligning against ChimeraSlayer Gold database provided with SILVA (Pruesse et al., 2007), second by using the *de novo* detection module in vsearch. *De novo* operational taxonomic unit (OTU) calling was performed on high-quality trimmed sequences at 97% similarity level using the USEARCH (Edgar, 2010) algorithm for clustering implemented in vsearch to generate operational taxonomical units (OTUs). Unique chimera filtered sequences were aligned using the Python Nearest Alignment Space Termination (PyNAST) (Caporaso et al., 2009) tool with a relaxed neighbour-joining tree built using FastTree (Price et al., 2010). The taxonomy was determined using the Classification Resources for Environmental Sequence Tags (CREST) (Lanzén et al., 2012) classifier with a confidence threshold of 0.80 against SILVA release 128 as a reference database.

Samples less than at least 2000 reads/sample were filtered from the OTU table in order to have sufficient reads as described in Caporaso et al. (2010a). After filtering, 386 samples were used for the statistical analyses, corresponding to 386 DNA extracts from 200 independent samples and ∼19.5 million reads (50,609 ± 26,700 reads/sample) assigned against 48,147 OTUs.

### Data Availability

The dataset is deposited at European Nucleotide Archive / SRA under the accession number PRJEB29109.

### Statistical Analysis

All statistical analyses were performed with a combination of QIIME1 V 1.90 (Caporaso et al., 2010b) and the R environment (Team, 2013). Multiple rarefactions were performed in QIIME with the smallest sample size as maximum depth. Alpha diversity (richness, Shannon and Simpson indices) were calculated in QIIME on the matrices resulting from the multiple rarefactions. The differences in alpha diversity indices were tested in QIIME using non-parametric (Monte-Carlo) tests across different pH categories with a Bonferroni correction.

The non-rarefied OTU-table was normalized using cumulative-sum scaling (CSS) in QIIME (Paulson et al., 2013). The resulting OTU-table was input into R for subsequent analyses. Using the phyloseq package (McMurdie and Holmes, 2013), we calculated the Bray-Curtis dissimilarity distance and visualized the ordination using principal coordinate analysis (PCoA). Using the cmdscale, envfit and ordisurf functions in the vegan package (Dixon, 2003), we calculated and visualized the significant correlations between edaphic properties and bacterial community dissimilarity in relation to the pH gradient. Permutational multivariate analyses of variance (PerMANOVA) was conducted using the adonis function with 999 permutations. Pearson’s correlation with 999 permutations was used to evaluate the collinearity of environmental variables.

To evaluate the spatial component, the geographic locations of the sampling sites were transformed into cartesian coordinates using the SoDA package (Chambers, 2008) and the euclidean distance was calculated using the vegan package. The presence of a linear trend was tested by redundancy discriminant analysis and ANOVA as prescribed in Borcard et al. (2018). A significant linear trend was identified, violating the second-order stationarity assumption where the mean of the variable and its spatial covariance are the same over the study area and its variance is finite. In other words, spatial correlation coefficients cannot be tested for significance if an overall trend is present in the data (Franklin and Mills, 2007; Borcard et al., 2018) and therefore, the data was detrended by linear regression of the x,y coordinates. To carry meaningful spatial analysis, constructing spatial variables representing spatial structures at all relevant scales is necessary (Borcard et al., 2018). To do so, we used distance-based Moran’s Eigenvector Maps (dbMEM) on detrended data and x,y coordinates using the adespatial R package (Dray et al., 2017). Significance of the spatial vectors was assessed using ANOVA. Forward selection was carried out to identify significant dbMEM vectors.

Variation partitioning analysis (VPA) was used to assess the impact of environmental and spatial factors on Bray-Curtis dissimilarity. The linear trend was considered to represent a source of variation like any other and therefore, we performed variation partitioning analysis using the environmental variables, x,y coordinates (trend) and significant dbMEM vectors.

Generalist and specialist taxa were identified by selecting taxa present in over 95 % of samples within each unique and overlapped pH category. Indicator taxa were determined by the Dufrene-Legendre indicator species analysis using the multipatt function (Cáceres and Legendre, 2009) to identify abundant OTUs that were specifically associated with the different pH ranges. The phylogenetic tree of indicator species was built using the aligned sequences from the identified indicator OTUs and input into FastTree. The tree was visualized using the Interactive Tree of Life (iTOL) online tool (Letunic and Bork, 2016).

## Results

### Bacterial community composition

We identified 48,147 bacterial taxa, mainly classified as Proteobacteria (20 %) and Planctomycetes (15%). Acidobacteria represented 13% while Actinobacteria, Bacteroidetes, Chloroflexi and Verrucomicrobia each accounted for 4 – 6 % of the community. Of these 48,147 OTUs, only 135 had overall abundances over 0.1% across all 386 samples and were defined as abundant taxa. They accounted for 32% of all the reads, illustrating the dominance of a few taxa over the rest of the community. Within these abundant taxa, Acidobacteria dominated the sub-community at approximately 32% with Blastocatellia (12.8%) and Acidobacteria subgroup6 (7.2%) as abundant classes. Alphaproteobacteria (10.6%) and Betaproteobacteria (6.6%) were the most commonly identified Proteobacteria (21% overall) and Verrucomicrobia was largely represented by Spartobacteria (17.2%). Actinobacteria (10.7%), Chloroflexi (7%) and Bacteroidetes (3.5%) were also among the abundant phyla classified.

### Spatial and environmental factors influencing bacterial communities

Distance-decay curves were used to evaluate the influence of spatial factors on bacterial community structure. Figure 2A illustrate the increase in community dissimilarity with increasing geographic distance (linear regression: R^2^ = 0.108, p *<*2.2 × 10^*-*16^). The best-fitted power model (R^2^ = 0.221, p = 0.005) suggested the presence of spatial autocorrelation, also qualified as dispersal limitation, within the first 500 m. For a more accurate estimate of dispersal limitation, we plotted the same graph within 0 and 500 m [Fig. 2B] and observed spatial autocorrelation approximately up to 20 m.

**Figure 2:**
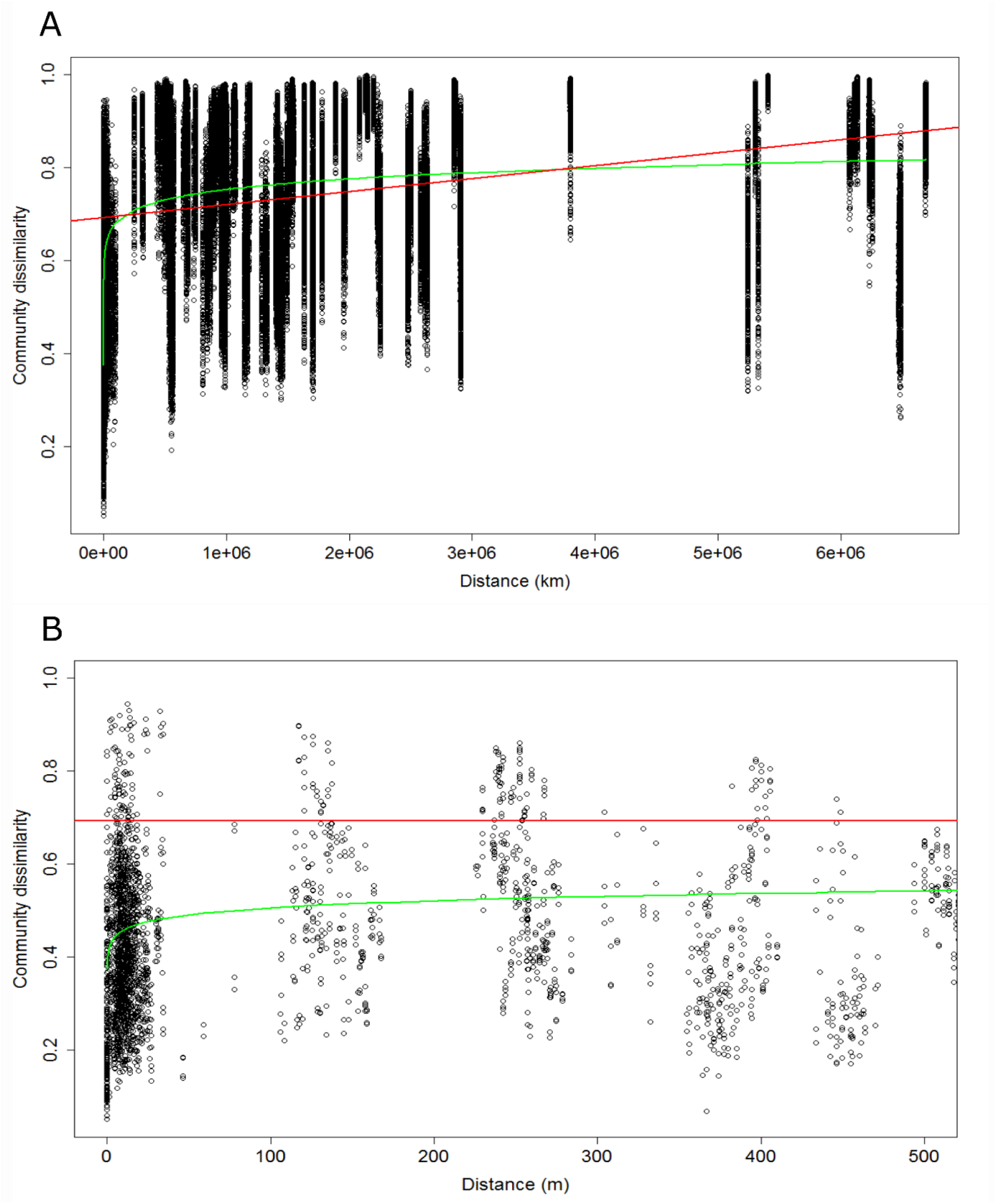
Distance-decay curves with the linear regression (red), the best-fitted power model (green). (A) Community dissimilarity of the total bacterial community by geographic distance in km. (B) Zoom into Figure 2A from 0 to 500 m to assess the spatial autocorrelation distance.

To determine the influence of environmental factors on the bacterial community structure, we used the adonis function with 999 permutations. Results indicated that pH explained 18.7 % of the variance in bacterial community composition overall [Table 1]. In comparison, conductivity, moisture and total organic carbon (TOC) explained almost 10 % of the variance all together. The measured environmental parameters explained 29 % of the variance in the bacterial sub-community, leaving 71 % unexplained. To quantify the influence of edaphic properties and spatial factors, we conducted variation partitioning on the bacterial community. We built spatial vectors using distance-based Moran’s Eigenvector Maps (dbMEM) on x,y geographic coordinates. Three vectors (MEMs) were produced; after forward selection, two significant MEMs were identified for the total community and used in further analyses. The variation partitioning analysis differentiated the effect of environmental factors, linear trend and spatial vectors on Bray-Curtis dissimilarity [Fig.3]. Results indicated that the environmental factors (X1 = [a]+[d]+[g]+[f]) explained 30.7 % of the variance in the community. These results are equivalent to the finding of the adonis test [Table 1], confirming the success of the variation partitioning analysis. Non-spatially structured environmental variables (fraction [a]) explained 15 % of the variation while spatial factors only (fractions [b], [e] and [c]) explained approximately 10 % of variation. Of fraction X1 (environmental factors), 15 % represented spatially structured environmental variables, also called induced spatial dependence, where the spatial structure of these environmental variables induced a similar spatial structure in the response data [detailed results in Table S1]. In total, 63.3 % of the total bacterial community dissimilarity could not be explained by environmental and spatial factors after R^2^ adjustment.

**Table 1:** The relative importance of environmental factors on bacterial communities, calculated by adonis.

| Variable | Total |  |
| --- | --- | --- |
|  | R<sup>2</sup> | Pr(>F) |
| pH | 0.187 | 0.0001*** |
| Conductivity | 0.048 | 0.0001*** |
| Moisture | 0.027 | 0.0001*** |
| TOC | 0.021 | 0.0001*** |
| Residuals | 0.717 | N/A |

**Figure 3:**
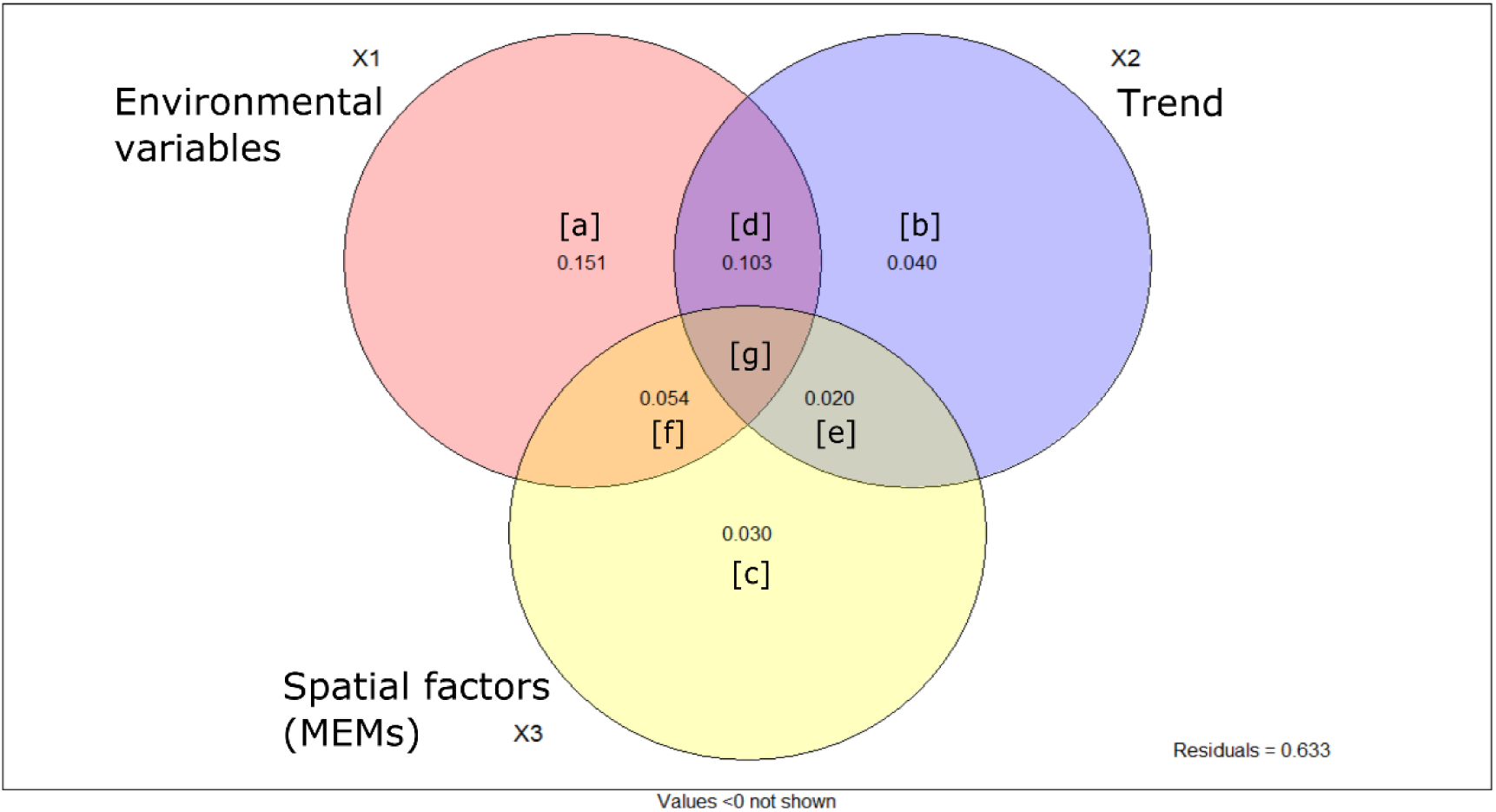
Venn diagram illustrating the results of the variation partitioning analysis and indicating the amount of variation in bacterial community explained by environmental variables and spatial factors including the linear trend. Detailed results are available in Table S1.

### pH as a key environmental factor

Of the environmental variables we measured, we identified pH as the primary factor structuring Arctic soil bacterial communities. The Pearson correlations indicated that pH did not have strong collinearity (coefficient over |0.8|) with any other variable measured [Table S2]. The clustering of samples by pH range was clearly observed on the principal coordinate analysis (PCoA) of bacterial communities [Fig. 4A]. The envfit analysis confirmed that pH was the key measured factor driving the ordination, although TOC, conductivity and moisture all had some influence. Using ordisurf, we visualised the PCoA ordination of the bacterial community in relation to the pH gradient [Fig. S1]. The Bray-Curtis dissimilarity heatmap and dendrogram [Fig. 4B] identified three main clusters illustrating community differences. The first cluster was composed of acidic samples in a pH gradient from Norwegian soils at pH = 4.07 (± 0.35) to samples from Alaska (pH = 4.64 ± 0.41) and West Greenland (pH = 4.90 ± 0.85). The second cluster included the lower acidoneutral range of samples from East Greenland (pH = 5.96 ± 0.69), Svalbard (pH = 5.65 ± 0.53) and Iceland (pH = 5.84 ± 0.46). Finally, the last cluster included the higher range of acidoneutral and alkaline soils from Russia (pH = 6.19 ± 0.22) and Canada (pH = 7.94 ± 0.67). The full spectrum of soil pH was covered from pH = 3.5 to pH = 9.0 and geographical locations often had samples from more than one pH category [Fig. S2]. As pH was the primary factor we identified influencing bacterial community structure, we defined three pH categories to focus on: acidic, acidoneutral and alkaline.

**Figure 4:**
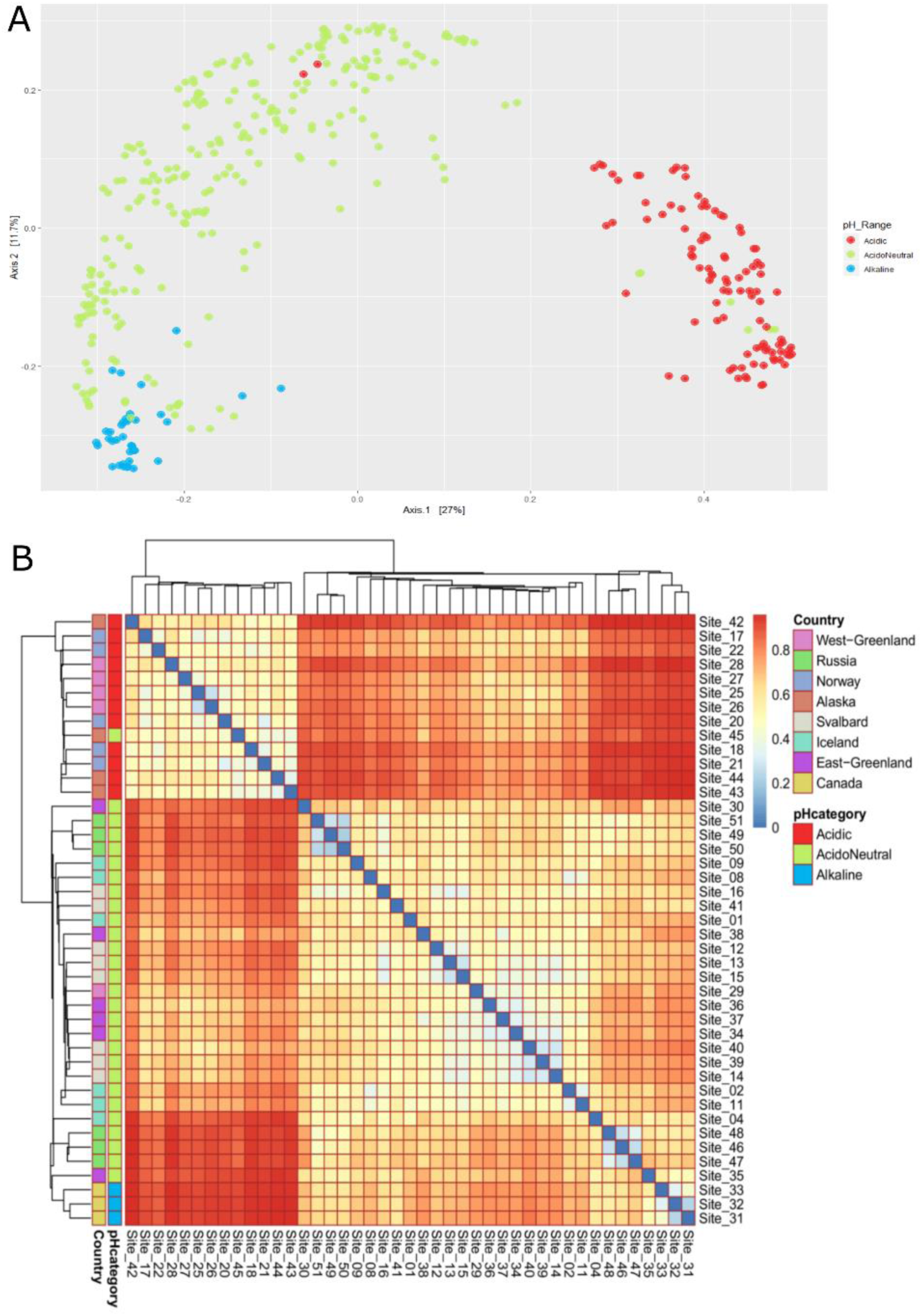
(A) PCoA of the Bray-Curtis dissimilarity of bacterial communities across the Arctic region by pH range. (B) Bray-Curtis dissimilarity heatmap and dendrogram illustrating the clustering of samples into three pH categories.

### Bacterial diversity by pH category

We compared the observed richness and Shannon diversity index by pH range [Fig. 5A, B] using a Monte-Carlo test to assess differences. Alpha-diversity was significantly lower in acidic samples than in acidoneutral and alkaline soils [Table S3]

**Figure 5:**
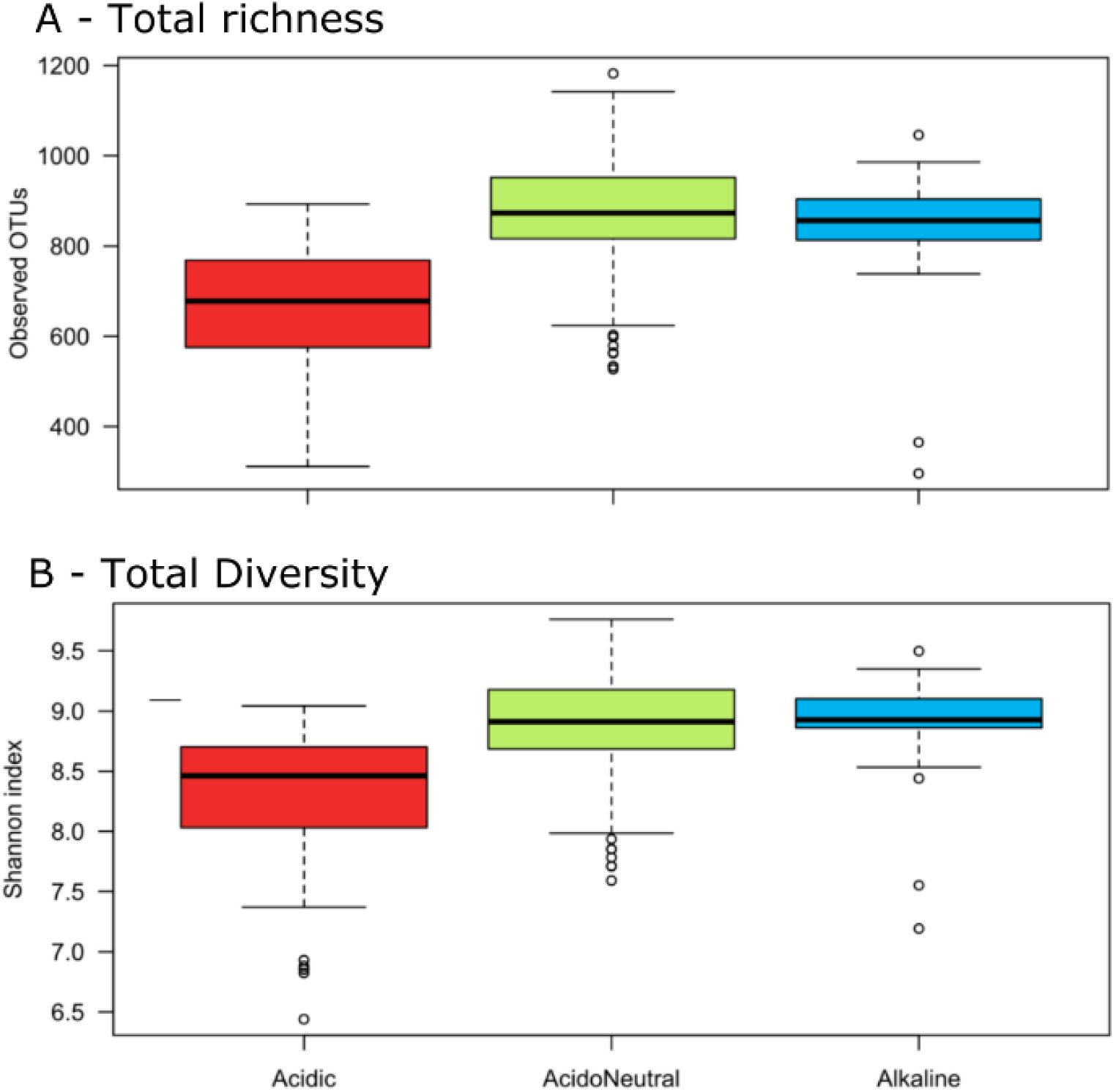
Alpha diversity differences in the total communities by pH category. (A) Richness by observed OTUs. (B) Shannon index.

In terms of community composition, we differentiated the abundant community, composed of only 135 OTUs, from the total community composed of 48,147 OTUs and observed the changes in relative abundance by pH range. In the total community, Acidobacteria, Armatimonadetes, Planctomycetes and Verrucomicrobia decreased with increasing pH. On the other hand, Bacteroidetes, Chloroflexi and Gemmatimonadetes increased in alkaline soils; Proteobacteria remained stable throughout. In the abundant community, Acidobacteria, Bacteroidetes, Gemmatimonadetes and SPAM increased with increasing pH while Actinobacteria, Planctomycetes and Verrucomicrobia remained stable [Fig. 6].

**Figure 6:**
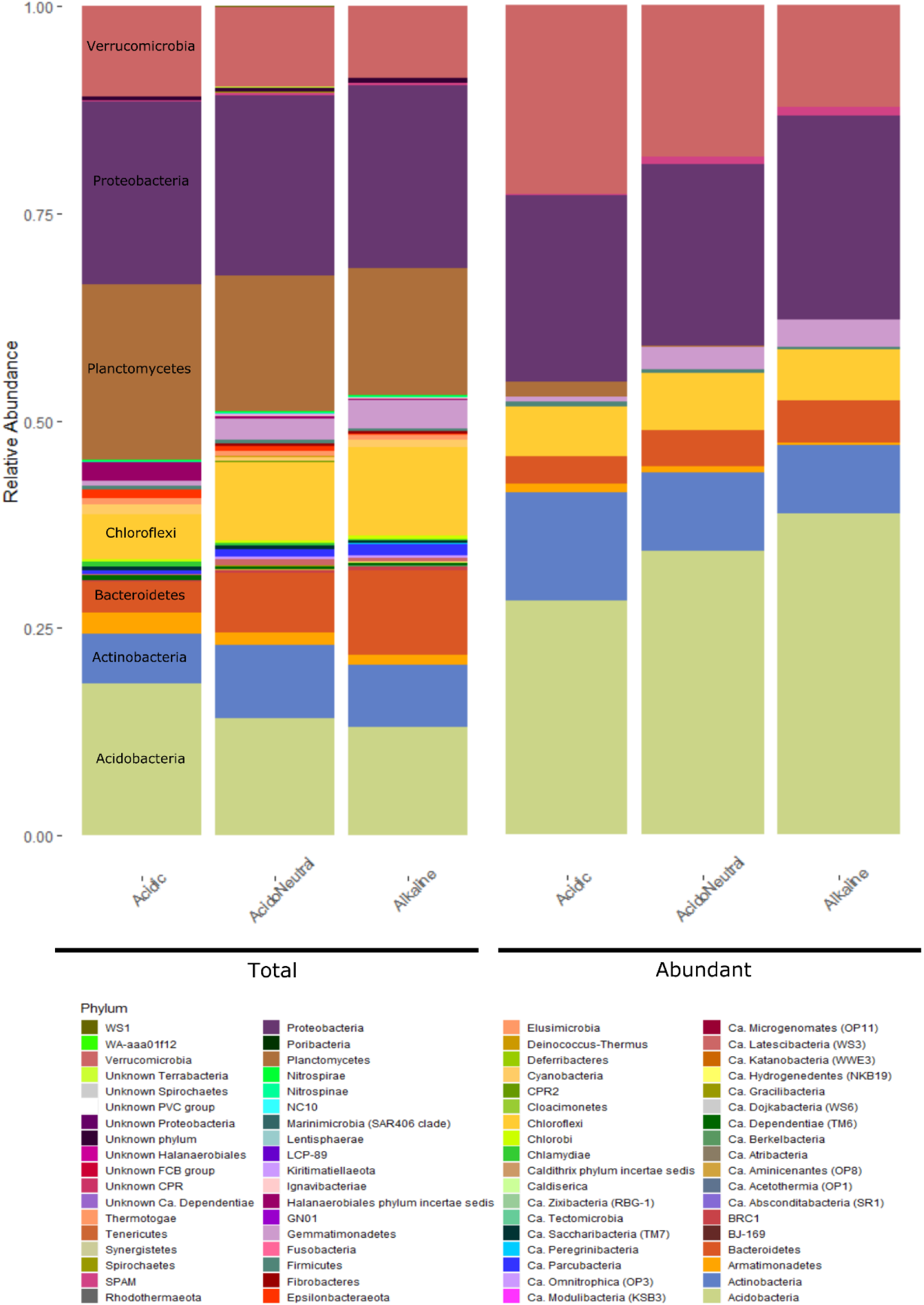
Relative abundance of the total and abundant communities by pH range classified at the phylum level.

### Generalist vs specialist taxa

The differentiation of specialist from generalist taxa associated with pH range was conducted by considering all taxa identified in this study and present in a minimum 95% of all samples from each pH category [Fig. 7]. 125 acidic specialist OTUs were identified and unique to acidic soil samples. Of these 125 unique OTUs, most belonged to the Acidobacteria (27%), Verrucomicrobia (20%), Actinobacteria (14%), Planctomycetes (14%) and Proteobacteria (14%). At the class level, Acidobacteria group 1 dominated at 14%, while the rest of the identified classes had balanced relative abundances oscillating between 5 and 8%. Shared OTUs, or generalists were present in low numbers in acidic soils. Only 12 OTUs were shared with acidoneutral soils only, which were dominated by Alphaproteobacteria (65%), mainly Rhizobiales, while 3 OTUs were shared with alkaline only, belonging to the Spartobacteria (Verrucomicrobia), Phycisphaerae (Planctomycetes) and Chitinophagia (Bacteroidetes) classes. In comparison, acidoneutral soils had 76 shared taxa with alkaline soils, against 12 shared OTUs with acidic. Taxa shared with alkaline soils were mainly classified as Blastocatellia (Acidobacteria), Spartobacteria and Acidobacteria subgroup 6. Taxa exclusively found in acidoneutral soils belonged mainly to Actinobacteria (20%), Verrucomicrobia (20%) and Acidobacteria (19%). At the class level, unique taxa living in acidoneutral soils were dominated by Spartobacteria, Alphaproteobacteria and Holophagae. Alkaline soils presented a combination of both, a high number of shared (79 OTUs in total) and 125 exclusive taxa. Alkaline unique taxa were mostly composed of Acidobacteria (36%) and Proteobacteria (22%). From these unique taxa, the Acidobacteria subgroup 6 (20%), Blastocatellia (12%) and Alphaproteobacteria (12%) dominated the community.

**Figure 7:**
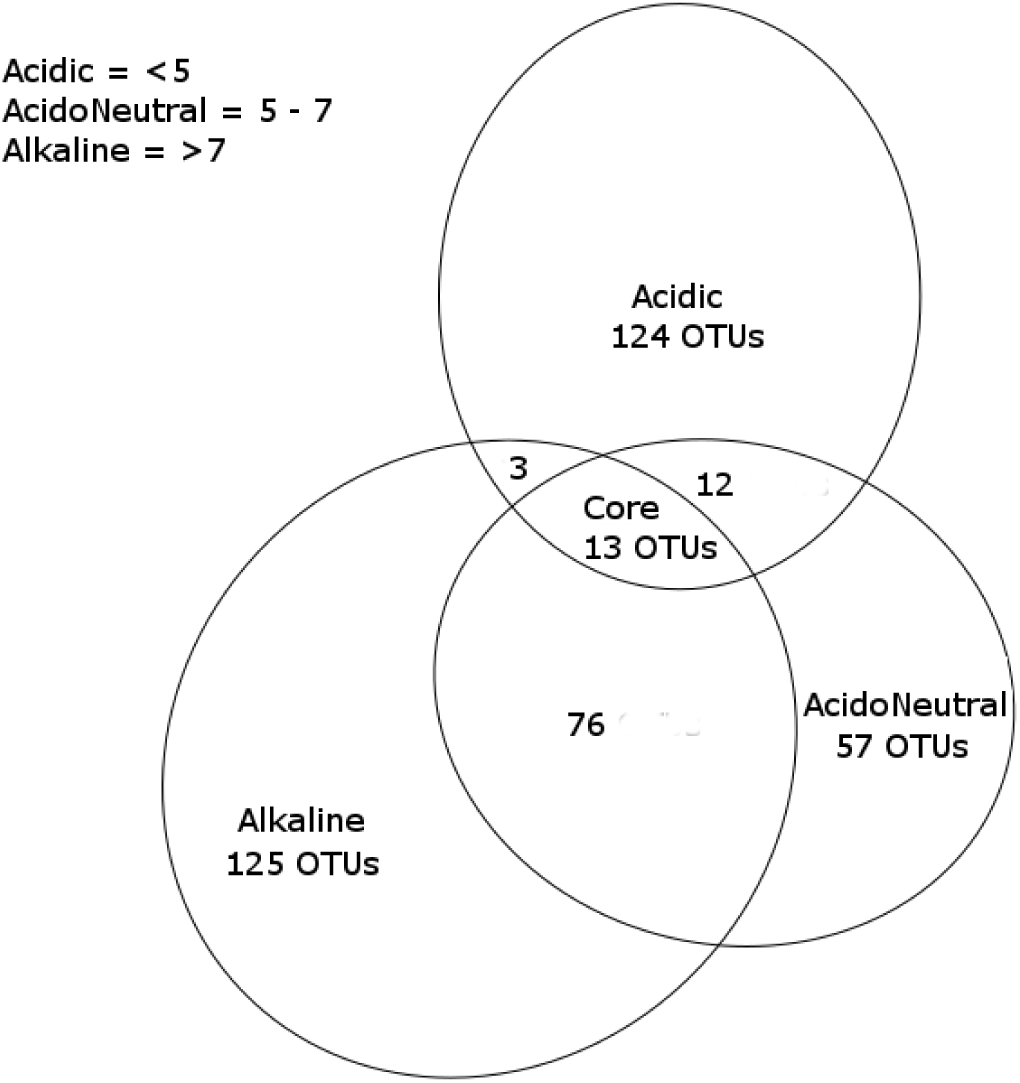
Venn diagram illustrating generalist (shared) and specialist (unique) OTUs by pH category.

### Indicator species

We conducted the indicator species analysis on the abundant taxa only as they may be more useful for monitoring purposes. We identified 17 unique taxa-habitat associations. In acidic soils, 10 indicator species, mainly Acidobacteria group 1 and 2 and Ca. Methylacidiphilum were identified. The 6 OTUs associated with Acidoneutral soils were mainly Acidobacteria group 4 (Blastocatellia) and Spartobacteria (Verrucomicrobia). Finally, only 1 OTU was identified as an indicator species for alkaline soils and belonged to the Holophagae (Acidobacteria). We also conducted the indicator species analysis of abundant taxa to combine pH ranges and identified 84 OTUs. 21 were identified as indicator species of acidic and acidoneutral soils combined, and mainly belonged to the Verrucomicrobia. Only 2 OTUs were associated with acidic and alkaline soils, Acidothermus (Actinobacteria) and Candidatus Xiphinematobacter (Verrucomicrobia), further illustrating the low overlap of taxa between these ecosystems and inferring the large ecosystem differences. Finally, 61 OTUs were associated with acidoneutral and alkaline soils, mainly belonging to the Acidobacteria and Proteobacteria phyla (see table S3 for the detailed list). The phylogenetic tree [Fig.S3] illustrates the classification of the identified abundant indicator taxa with the pH range associated. Although displaying diversity, they were mainly classified as Acidobacteria and Verrucomicrobia, with the vast majority of taxa associated with acidoneutral and alkaline soils.

### The core microbiome

While some taxa displayed biogeographical patterns with non-random distribution across space and environmental gradients, others were cosmopolitan and identified in over 95 % of all samples. The core microbiome represented 0.026 % of bacterial communities and accounted for 2.7 % of all reads. It was composed of 13 OTUs, mainly Proteobacteria and Acidobacteria, notably belonging to the Rhizobiales and Acidobacteria subgroup 6. The Alphaproteobacteria, especially the Bradyrhizobiaceae family was most abundant in acidic soils and decreased with increasing pH. The Betaproteobacteria, especially the Comamonadaceae family, decreased with decreasing pH. The Acidobacteria were highest in the acidoneutral soils. Although relative abundances differed by pH range, these 13 OTUs were present in all categories. Figure S4 illustrates the taxonomic classes of the core microbiome and distribution between each pH category.

## Discussion

The coverage of this study allowed the identification of a core microbiome, present in over 95 % of the samples analyzed and composed of only 13 OTUs. The most abundant of these taxa belonged to the Bradyrhizobiaceae family. This is one of the most common families worldwide, as identified by Delgado-Baquerizo et al. (2018). However, whether this is the same taxa remains unclear and highlights the need for global studies incorporating extreme environments. The identification of a core Arctic soil microbiome is novel and this low number of cosmopolitan OTUs illustrates the potential for endemism. For the rest of the bacterial community, spatial and edaphic factors influenced distribution across the region.

### Arctic bacterial community assembly

The distance-decay curve [Fig.2A,2B] illustrated the influence of geography on bacterial communities. The spatial autocorrelation is a proxy for dispersal limitation, which was restricted to approximately 20 m [Fig.2B]. However, the lack of small-scale data is problematic in estimating the real autocorrelation range. The variation partitioning analysis quantified the importance of both, selection (deterministic) and dispersal (stochastic) on the bacterial community structure. Environmental variables explained 30 % of the total variation (selection), of which 15 % was spatially structured, corresponding to the induced spatial dependence, as described in Borcard et al. (2018). Spatial components (trend + MEMs) alone explained 9 % of the variation, illustrating the spatial autocorrelation (Borcard et al., 2018), or dispersal.

Here, we focused on environmental variables previously identified as influencing bacterial communities elsewhere and included pH, conductivity, moisture and organic carbon. However, 63 % of the variation remained unexplained by either environmental or spatial factors in the variation partitioning analysis. This unexplained variance could be the result of unmeasured biotic or abiotic factors, directly or indirectly influencing bacterial communities. Examples of unmeasured edaphic properties could include total nitrogen, zinc, calcium oxide or nickel, all shown to have an influence on bacterial community structure in the Arctic (Wojcik et al., 2018). Other ecosystem properties such as ice presence, active layer depth or soil texture are likely to have some influence. Climatic and topographic variables such as temperature, precipitation (Yergeau and Kowalchuk, 2008; Castro et al., 2010; Nielsen and Ball, 2015) and altitude (Shen et al., 2013; Xu et al., 2014) are also known to influence bacterial assemblage and were not included in this study. Vegetation cover is a known driver of bacterial community assemblage in the Arctic region (as reviewed by Malard and Pearce (2018)) which was not included in this analysis. Biotic interactions may also be a major part of the unexplained variance observed and could include competition and predation within bacterial communities or with other members of the soil biota (Wardle, 2006; Singh et al., 2009). Other processes influencing bacterial communities include diversification and drift, both likely to account for some of the unexplained variance (Martiny et al., 2006).

### The importance of pH

We evaluated the impact of environmental variables known to influence global soil bacterial communities. Overall, pH explained the most variation (18.7 %) in bacterial community structure and diversity while TOC, moisture and conductivity explained between 2 and 5 % of the variance respectively. The identification of pH as a key factor influencing Arctic soil bacterial community composition is in line with previous Arctic studies over both, small and large scales (Männistö et al., 2006; Chu et al., 2010; Ganzert et al., 2014; Siciliano et al., 2014; Schostag et al., 2015). However, this study is the first to quantify the magnitude of influence, highlighting the large variation in bacterial community remaining unexplained. While pH has been identified globally as a major factor influencing microbial diversity and community structure (Fierer and Jackson, 2006; Lauber et al., 2009; Tedersoo et al., 2014; Delgado-Baquerizo et al., 2018), the underlying processes and mechanisms by which it does remain unclear. Studies have demonstrated that the soil pH is correlated with other elements of the geochemistry and has a strong impact on nutrient and water availability as well as solubility and adsorption (Gray et al.,2014). For instance, acidic pH increases aluminium, hydrogen and manganese solubility, retarding plant root growth due to high toxicity (Clark and Baligar, 2000; Singh et al., 2017). Acidic soils also have nutrient deficiencies such as calcium magnesium and potassium but also decreased phosphorus and molybdenum solubilities (Baligar and Clark, 2000; Gray et al., 2014). Alkaline soils are generally the result of low precipitation and high evapotranspiration, leading to low water availability and in common with acidic soils, nutrient deficiencies are found with decreased phosphorus, iron, copper or zinc for instance (Baligar and Clark, 2000). In similar ways, acidic and alkaline soils are generally considered harsh environments requiring a wide range of adaptations from microorganisms while acidoneutral soils are considered the optimum environment for microbial life (Fierer and Jackson, 2006; Rousk et al., 2010); such differences in soil composition are likely driving the observed differences in microbial community composition by pH range.

### Distribution of generalist and specialist taxa

Microbial communities are assembled by deterministic (selection) and stochastic (dispersal) processes. It has been hypothesized that communities primarily structured by deterministic processes will host more specialist taxa, highly adapted to the ecosystem, while communities influenced by dispersal will harbour primarily generalist taxa, more resilient to change (Pandit et al., 2009; Graham and Stegen, 2017; Sriswasdi et al., 2017). While specialist taxa are restricted to certain niches, they can be locally abundant; shared taxa, or generalists, are distributed across many niches (Barberán et al., 2012). In most cases specialist will be more abundant because generalists rapidly become specialists to adapt to their ecosystems, despite generalists having evolutionary advantages (Sriswasdi et al., 2017).

In this study, bacterial communities were specialist-dominated in acidic soils, generalist-dominated in acidoneutral soils and a mixed community in alkaline samples. The higher abundance of specialists in acidic soils (considered the harshest systems) could illustrate the need for environmental adaptations to survive in these ecosystems and suggests that deterministic processes likely structure microbial communities. Geographically, the first cluster identified by the Bray-Curtis dissimilarity matrix [Fig. 4B] grouped all acidic samples from northern Norway, western Greenland and Alaska, illustrating the similarities of their bacterial communities despite the large distances separating these locations, further suggesting the strong influence of selection over dispersal. The dominance of generalists in acidoneutral soils illustrates the lower environmental pressure to have specific survival adaptations and infers the dominance of stochastic processes in community structuring. Eastern Greenland, Svalbard and Iceland were grouped together by the Bray-Curtis dissimilarity matrix. Geographically, these locations connected through the Greenland sea, a possible route of dispersal through aerosolization and long-distance aeolian transport (Šantl-Temkiv et al., 2018). Finally, alkaline soils hosted a mixed-community of specialists and generalists, suggesting the combination of both, selection and dispersal, in community structuring. The abundance of generalist taxa in alkaline soils, generally considered a harsh system in many ways similar to acidic soils, highlights the adaptability of generalists to a wide range of environmental conditions. Interestingly, the Bray-Curtis dissimilarity matrix clustered the Canadian (alkaline) and Russian (acidoneutral) samples together. The Russian samples were on the higher end of the acidoneutral pH scale and the grouping of these samples illustrates the blurred boundary between acidoneutral and alkaline pH categories. While environmental selection may be driven by many variables, dispersal likely occurs via long-distance aeolian dispersal (Šantl-Temkiv et al., 2018).

### The importance of indicator taxa

The indicator species analysis determined OTU-pH range associations identifying different classes of Acidobacteria and Verrucomicrobia, primarily, characteristic of each pH range. The identification of abundant indicator species with strong habitat associations opens the possibility of predicting the presence and abundance of these taxa across the Arctic region. This is especially important for taxa such as Ca. Methylacidiphilum, a known methanotroph unlike others as it belongs to the Verrucomicrobia phylum instead of the Proteobacteria (Dunfield et al., 2007; Khadem et al., 2010). While natural CH_4_ emissions come primarily from wetlands, the identification of these phylotypes in acidic soils suggests wetland may not be the only major source of northern methane emissions. Understanding the distribution and abundance of such indicator taxa, combined with field gas measurements may allow large-scale estimates of climate-active gases release rates in the region, from soils and wetlands (Wartiainen et al., 2006; Jørgensen et al., 2015).

### Concluding remarks

This study investigated patterns in soil bacterial diversity across the Arctic and demonstrated that bacterial communities differed across the region. However, the dispersal range was limited to approximately 20 m and the variation in communities across the Arctic was primarily driven by environmental rather than spatial factors. The key edaphic property influencing communities we identified was pH, in accordance with that seen elsewhere. However, 63 % of the variation remained unexplained, calling for more in-depth investigations of the drivers of bacterial community structure in the region. We identified unique and indicator taxa closely associated to the soil pH. Finally, a core microbiome composed of only 13 ubiquitously distributed taxa was identified, highlighting the potential for endemism in the region. While this study brings a deeper understanding of Arctic bacterial community assemblages, this is also a baseline for future functional studies in the region, which will be critical to forecast the ecological consequences of environmental change.

## Acknowledgements

This work was supported by a grant from the European Commission’s Marie Sklowdowska Curie Actions program under project number 675546. The authors also thank Dr Paul Mann, Dr Charles Greer, Dr Svetlana Evgrafova, Dr Bill Holben, Sam Pannoni, Edwin Sia and UNIS for their participation and support in fieldwork. MiSeq sequencing of the 16S rRNA gene was performed by the NU-OMICS sequencing service (Northumbria University).

