## Supplementary material for "Biogeographical patterns in soil bacterial communities across the Arctic region"

### Supplementary Figures

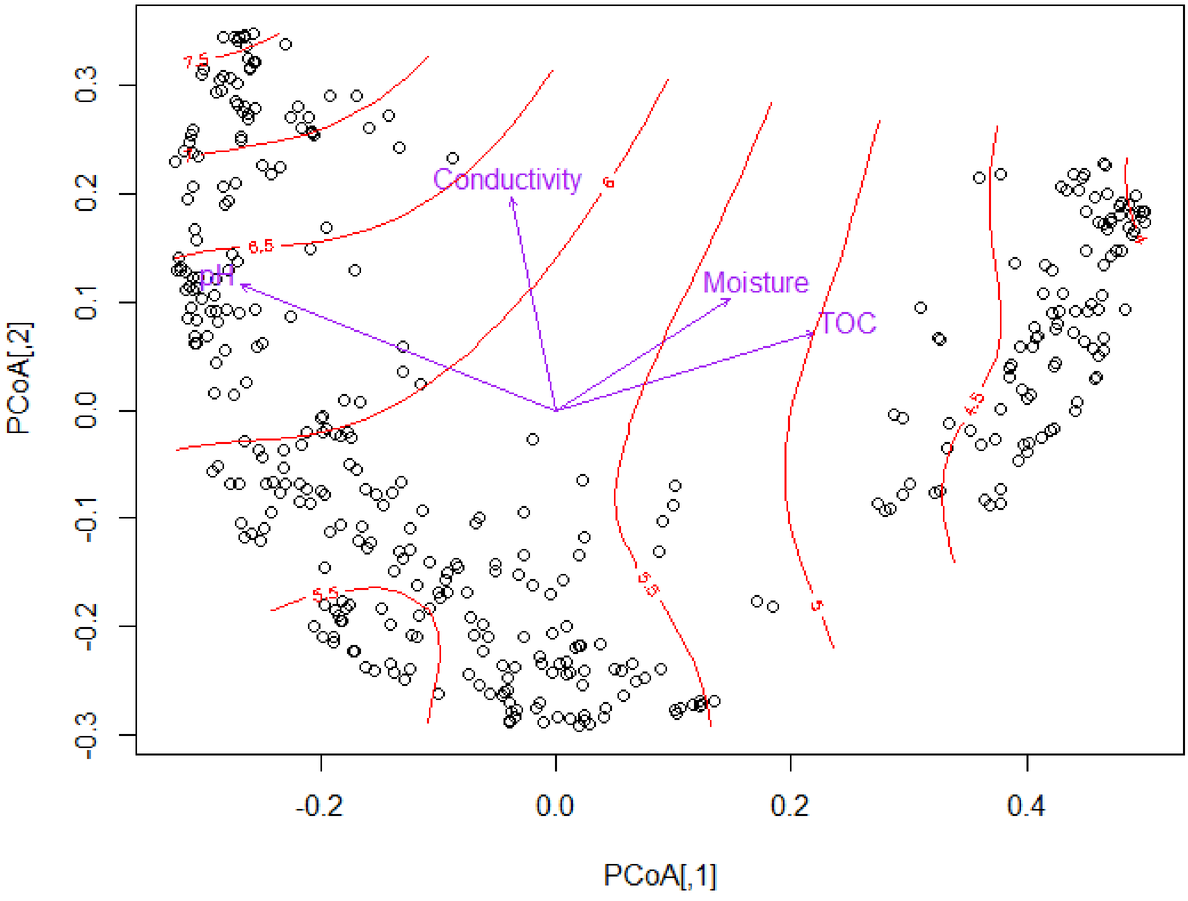

*Fig S1: PCoA of Bray-Curtis dissimilarity with the pH gradient in red and the vectors illustrating the influence of environmental variables on the ordination.*

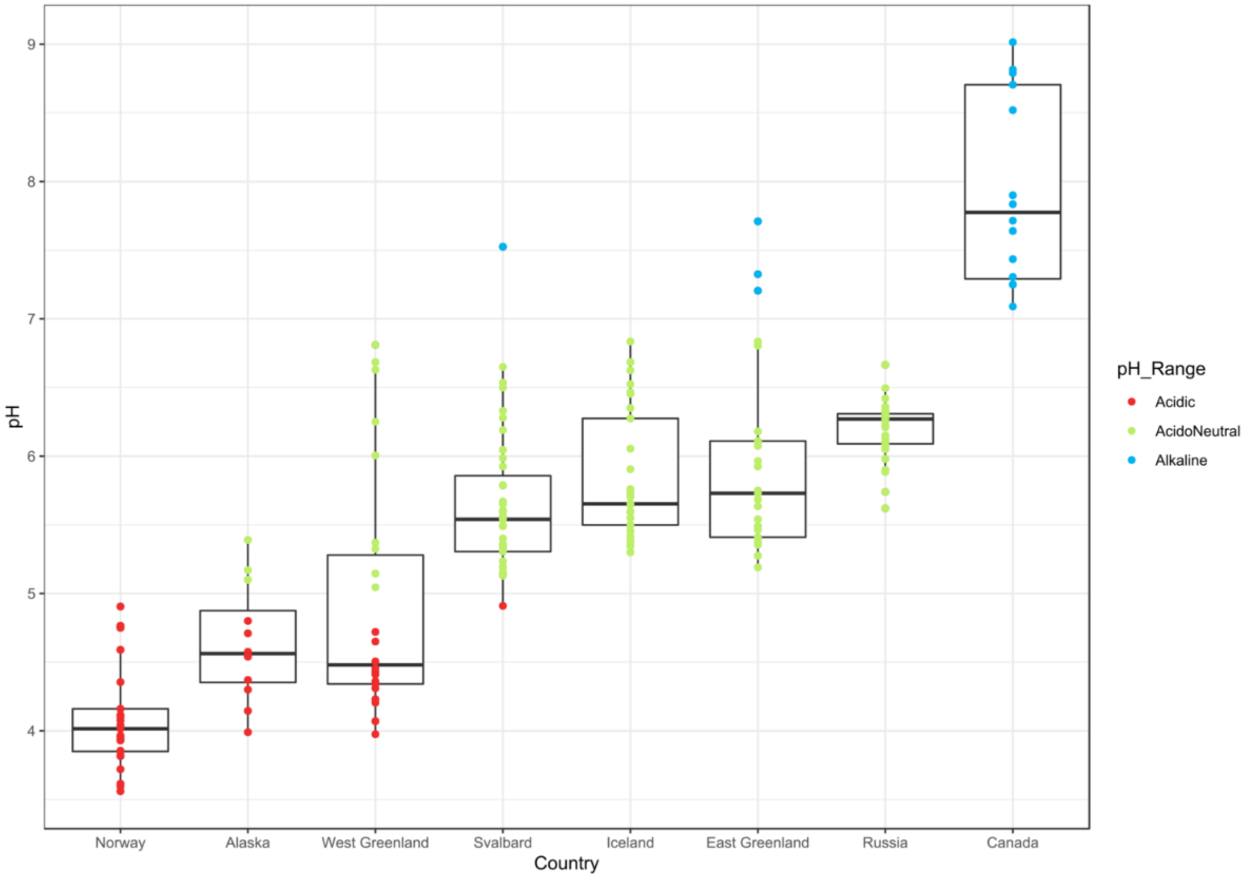

*Fig S2: pH gradient sampling curve illustrating the broad coverage of the pH gradient and the pH variability within locations.*

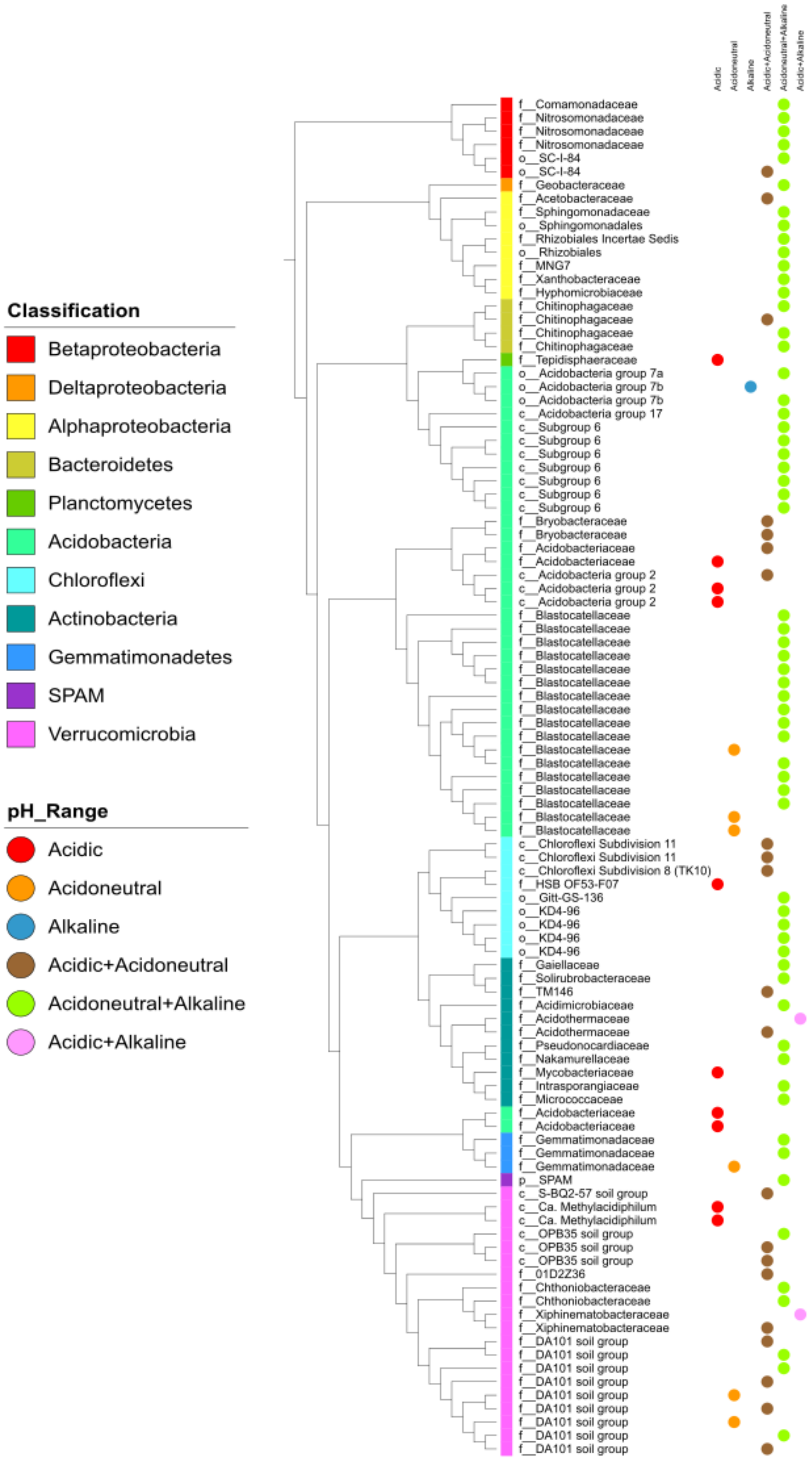

*Fig S3: Phylogenetic tree of abundant indicator taxa associated to a pH category.*

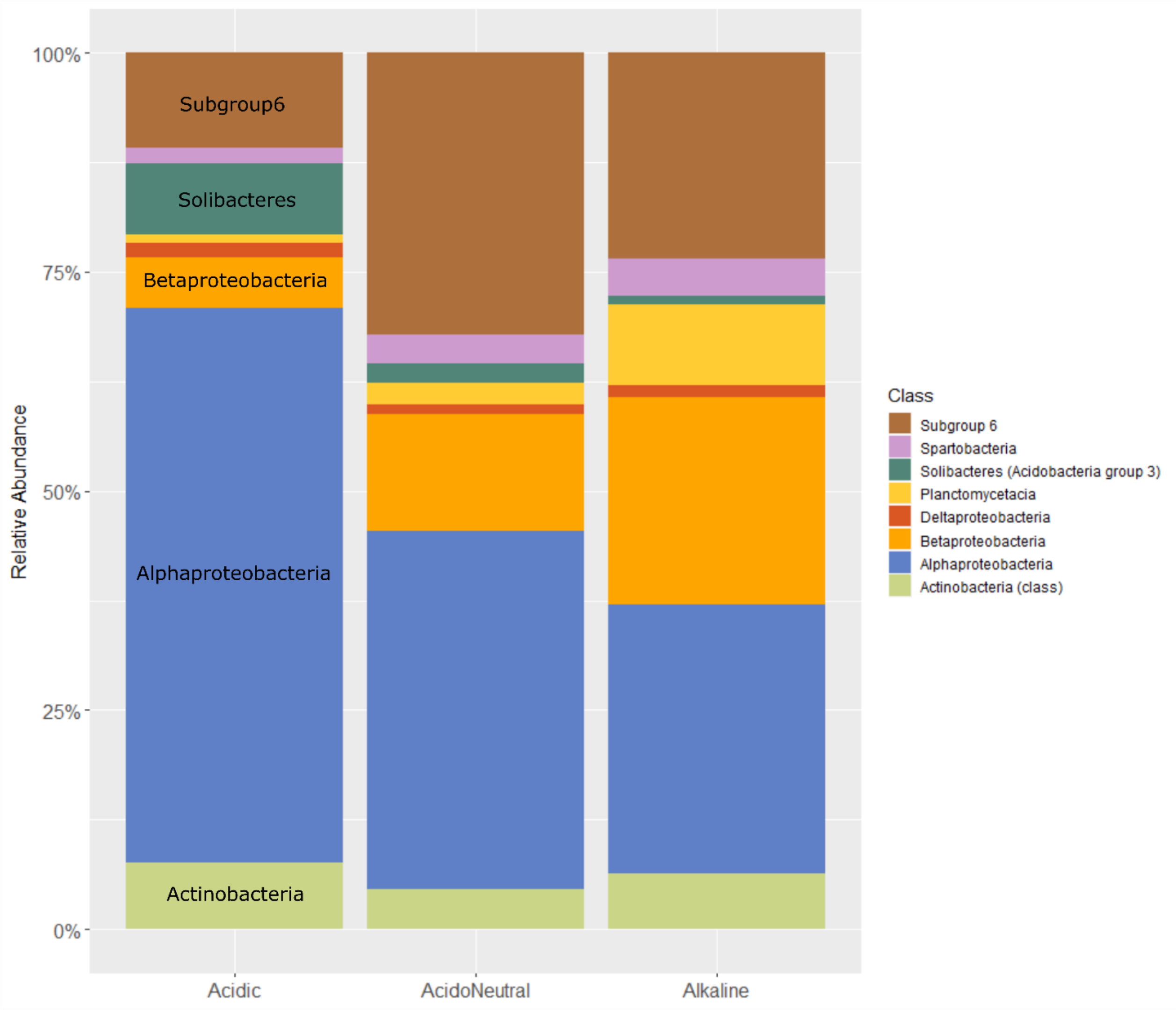

*Figure S4: Relative abundance of the Arctic core microbiome by pH range classified at the class level.*

### Supplementary Tables

*Table S1: Results of the variation partitioning analysis. Partition X1 represents the environmental variables, partition X2 represents the linear trend while partition X3 represents the spatial vectors (MEMs). Detailed fractions are illustrated in Figure 3 of the main document.*

| | Total | | |
|---|---|---|---|
| **Partition table** | **Df** | **$R^2$** | **Adj. $R^2$** |
| [a+d+f+g] = X1 | 4 | 0.284 | 0.276 |
| [b+d+e+g] = X2 | 2 | 0.136 | 0.131 |
| [c+e+f+g] = X3 | 2 | 0.077 | 0.073 |
| [a+b+d+e+f+g] = X1+X2 | 6 | 0.347 | 0.337 |
| [a+c+d+e+f+g] = X1+X3 | 6 | 0.337 | 0.327 |
| [b+c+d+e+f+g] = X2+X3 | 4 | 0.224 | 0.216 |
| [a+b+c+d+e+f+g] = All | 8 | 0.380 | 0.367 |
| **Individual fractions** | **Df** | **$R^2$** | **Adj. $R^2$** |
| [a] = X1 / X2+X3 | 4 | N/A | 0.151 |
| [b] = X2 / X1+X3 | 2 | N/A | 0.04 |
| [c] = X3 / X1+X2 | 2 | N/A | 0.03 |
| [d] = X1+X2 | 0 | N/A | 0.103 |
| [e] = X1+X3 | 0 | N/A | 0.02 |
| [f] = X2+X3 | 0 | N/A | 0.054 |
| [g] = All | 0 | N/A | -0.032 |
| [h] = Residuals | N/A | N/A | 0.633 |

*Table S2: Pearson correlations physiochemical properties using Multivariate Data Analysis.*

| | pH | Conductivity | Moisture | TOC |
|---|---|---|---|---|
| pH | 1.000 | 0.394 | -0.442 | -0.556 |
| Conductivity | 0.394 | 1.000 | 0.040 | 0.099 |
| Moisture | -0.442 | 0.040 | 1.000 | 0.829 |
| TOC | -0.556 | 0.099 | 0.829 | 1.000 |

*Table S3: Non-parametric (Monte Carlo) test on the observed richness and Shannon index across different pH catego ries with Bonferroni correction.*

Observed species statistical comparison

| Group1 | Group2 | Group1 mean | Group1 std | Group2 mean | Group2 std | t stat | p-value |
|---|---|---|---|---|---|---|---|
| Acidic | AcidoNeutral | 660.318 | 134.705 | 871.094 | 113.337 | -14.823 | 0.003 |
| Acidic | Alkaline | 660.318 | 134.705 | 838.325 | 137.801 | -6.708 | 0.003 |
| Alkaline | AcidoNeutral | 838.325 | 137.801 | 871.094 | 113.337 | -1.569 | 0.333 |

Shannon index statistical comparison

| Group1 | Group2 | Group1 mean | Group1 std | Group2 mean | Group2 std | t stat | p-value |
|---|---|---|---|---|---|---|---|
| Acidic | AcidoNeutral | 8.309 | 0.558 | 8.889 | 0.386 | -11.065 | 0.003 |
| Acidic | Alkaline | 8.309 | 0.558 | 8.889 | 0.430 | -5.621 | 0.003 |
| Akaline | AcidoNeutral | 8.889 | 0.430 | 8.889 | 0.386 | 0.0003 | 1.0 |

*Table S4: Indicator species determined by the Dufrene-Legendre indicator species analysis method to identify abundant OTUs (>0.1%) habitat-associated with the different pH ranges.*

```
Multilevel pattern analysis
 ---------------------------

 Association function: IndVal.g
 Significance level (alpha): 0.05

 Total number of OTUs: 134
 Selected number of OTUs: 101
 Number of OTUs associated to 1 group: 17
 Number of OTUs associated to 2 groups: 84
```

| pH Category | OTUs * | Indicator value | P value | Taxonomy ** |
|---|---|---|---|---|
| Acidic | OTU_115 | 0.999 | 0.001 | k__Bacteria; p__Acidobacteria; c__Acidobacteria group 2; |
| | OTU_57 | 0.998 | 0.001 | k__Bacteria; p__Acidobacteria; c__Acidobacteria group 2; |
| | OTU_88 | 0.998 | 0.001 | k__Bacteria; p__Actinobacteria; c__Actinobacteria (class); o__Corynebacteriales; f__Mycobacteriaceae; g__Mycobacterium; |
| | OTU_160 | 0.995 | 0.001 | k__Bacteria; p__Planctomycetes; c__Phycisphaerae; o__Tepidisphaerales; f__Tepidisphaeraceae; |
| | OTU_41 | 0.977 | 0.001 | k__Bacteria; p__Verrucomicrobia; c__Ca. Methylacidiphilum; |
| | OTU_625 | 0.969 | 0.002 | k__Bacteria; p__Verrucomicrobia; c__Ca. Methylacidiphilum; |
| | OTU_73 | 0.963 | 0.004 | k__Bacteria; p__Acidobacteria; c__Acidobacteriia (Acidobacteria group 1); o__Acidobacteriales; f__Acidobacteriaceae; |
| | OTU_48 | 0.961 | 0.001 | k__Bacteria; p__Acidobacteria; c__Acidobacteriia (Acidobacteria group 1); o__Acidobacteriales; f__Acidobacteriaceae; |
| | OTU_471 | 0.954 | 0.001 | k__Bacteria; p__Acidobacteria; c__Acidobacteriia (Acidobacteria group 1); o__Acidobacteriales; f__Acidobacteriaceae; |
| | OTU_181 | 0.917 | 0.025 | k__Bacteria; p__Chloroflexi; c__Ktedonobacteria; o__Ktedonobacterales; f__HSB OF53-F07; |
| AcidoNeutral | OTU_25190 | 0.999 | 0.001 | k__Bacteria; p__Acidobacteria; c__Blastocatellia (Acidobacteria group 4); o__Blastocatellales; f__Blastocatellaceae; g__RB41; |
| | OTU_13780 | 0.995 | 0.001 | k__Bacteria; p__Acidobacteria; c__Blastocatellia (Acidobacteria group 4); o__Blastocatellales; f__Blastocatellaceae; g__RB41; |

| | | | | |
|---|---|---|---|---|
| | OTU_9640 | 0.993 | 0.001 | k__Bacteria; p__Verrucomicrobia; c__Spartobacteria; o__Chthoniobacterales; f__DA101 soil group; |
| | OTU_47944 | 0.993 | 0.001 | k__Bacteria; p__Acidobacteria; c__Blastocatellia (Acidobacteria group 4); o__Blastocatellales; f__Blastocatellaceae; g__RB41; |
| | OTU_23501 | 0.976 | 0.001 | k__Bacteria; p__Gemmatimonadetes; c__Gemmatimonadetes (class); o__Gemmatimonadales; f__Gemmatimonadaceae; |
| | OTU_46058 | 0.975 | 0.001 | k__Bacteria; p__Verrucomicrobia; c__Spartobacteria; o__Chthoniobacterales; f__DA101 soil group; |
| Alkaline | OTU_101 | 0.931 | 0.001 | k__Bacteria; p__Acidobacteria; c__Holophagae; o__Acidobacteria group 7b; |
| Acidic + AcidoNeutral | OTU_15 | 1 | 0.001 | k__Bacteria; p__Proteobacteria; c__Betaproteobacteria; o__SC-I-84; |
| | OTU_22 | 1 | 0.001 | k__Bacteria; p__Acidobacteria; c__Solibacteres (Acidobacteria group 3); o__Solibacterales; f__Bryobacteraceae; g__Bryobacter; |
| | OTU_137 | 1 | 0.001 | k__Bacteria; p__Proteobacteria; c__Alphaproteobacteria; o__Rhodospirillales; f__Acetobacteraceae; |
| | OTU_35 | 1 | 0.001 | k__Bacteria; p__Bacteroidetes; c__Chitinophagia; o__Chitinophagales; f__Chitinophagaceae; |
| | OTU_71 | 0.987 | 0.001 | k__Bacteria; p__Verrucomicrobia; c__OPB35 soil group; |
| | OTU_29 | 0.987 | 0.001 | k__Bacteria; p__Verrucomicrobia; c__OPB35 soil group; |
| | OTU_121 | 0.987 | 0.001 | k__Bacteria; p__Verrucomicrobia; c__S-BQ2-57 soil group; |
| | OTU_85 | 0.987 | 0.001 | k__Bacteria; p__Actinobacteria; c__Thermoleophilia; o__Solirubrobacterales; f__TM146; |
| | OTU_79 | 0.975 | 0.002 | k__Bacteria; p__Verrucomicrobia; c__Spartobacteria; o__Chthoniobacterales; f__Xiphinematobacteraceae; g__Candidatus Xiphinematobacter; |
| | OTU_28 | 0.975 | 0.002 | k__Bacteria; p__Acidobacteria; c__Acidobacteria group 2; |
| | OTU_53 | 0.975 | 0.004 | k__Bacteria; p__Chloroflexi; c__Chloroflexi Subdivision 8 (TK10); |
| | OTU_212 | 0.975 | 0.001 | k__Bacteria; p__Verrucomicrobia; c__Spartobacteria; o__Chthoniobacterales; f__01D2Z36; |
| | OTU_14 | 0.975 | 0.005 | k__Bacteria; p__Chloroflexi; c__Chloroflexi Subdivision 11; |

| | | | | |
|---|---|---|---|---|
| | OTU_62 | 0.975 | 0.003 | k__Bacteria; p__Acidobacteria; c__Acidobacteriia (Acidobacteria group 1); o__Acidobacteriales; f__Acidobacteriaceae; |
| | OTU_30300 | 0.974 | 0.012 | k__Bacteria; p__Verrucomicrobia; c__Spartobacteria; o__Chthoniobacterales; f__DA101 soil group; |
| | OTU_164 | 0.962 | 0.003 | k__Bacteria; p__Verrucomicrobia; c__Spartobacteria; o__Chthoniobacterales; f__DA101 soil group; |
| | OTU_108 | 0.962 | 0.007 | k__Bacteria; p__Chloroflexi; c__Chloroflexi Subdivision 11; |
| | OTU_408 | 0.962 | 0.011 | k__Bacteria; p__Verrucomicrobia; c__Spartobacteria; o__Chthoniobacterales; f__DA101 soil group; |
| | OTU_65 | 0.948 | 0.013 | k__Bacteria; p__Actinobacteria; c__Actinobacteria (class); o__Frankiales; f__Acidothermaceae; g__Acidothermus; |
| | OTU_112 | 0.922 | 0.011 | k__Bacteria; p__Verrucomicrobia; c__Spartobacteria; o__Chthoniobacterales; f__DA101 soil group; |
| | OTU_78 | 0.908 | 0.02 | k__Bacteria; p__Acidobacteria; c__Solibacteres (Acidobacteria group 3); o__Solibacterales; f__Bryobacteraceae; g__Bryobacter; |
| Acidic + Alkaline | OTU_39 | 0.992 | 0.001 | k__Bacteria; p__Actinobacteria; c__Actinobacteria (class); o__Frankiales; f__Acidothermaceae; g__Acidothermus; |
| | OTU_23 | 0.957 | 0.02 | k__Bacteria; p__Verrucomicrobia; c__Spartobacteria; o__Chthoniobacterales; f__Xiphinematobacteraceae; g__Candidatus Xiphinematobacter; |
| AcidoNeutral + Alkaline | OTU_15656 | 1 | 0.001 | k__Bacteria; p__Acidobacteria; c__Blastocatellia (Acidobacteria group 4); o__Blastocatellales; f__Blastocatellaceae; g__RB41; |
| | OTU_52 | 1 | 0.001 | k__Bacteria; p__Acidobacteria; c__Subgroup 6; |
| | OTU_96 | 1 | 0.001 | k__Bacteria; p__Proteobacteria; c__Alphaproteobacteria; o__Sphingomonadales; f__Sphingomonadaceae; g__Sphingomonas; |
| | OTU_155 | 1 | 0.001 | k__Bacteria; p__Acidobacteria; c__Subgroup 6; |
| | OTU_11936 | 1 | 0.001 | k__Bacteria; p__Acidobacteria; c__Blastocatellia (Acidobacteria group 4); o__Blastocatellales; f__Blastocatellaceae; g__RB41; |
| | OTU_26 | 1 | 0.001 | k__Bacteria; p__Acidobacteria; c__Blastocatellia (Acidobacteria group 4); o__Blastocatellales; f__Blastocatellaceae; |
| | OTU_67 | 1 | 0.001 | k__Bacteria; p__Acidobacteria; c__Blastocatellia (Acidobacteria group 4); o__Blastocatellales; f__Blastocatellaceae; |

| | | | | |
|---|---|---|---|---|
| | OTU_81 | 1 | 0.001 | k__Bacteria; p__Proteobacteria; c__Alphaproteobacteria; o__Sphingomonadales; |
| | OTU_1779 | 1 | 0.001 | k__Bacteria; p__Acidobacteria; c__Blastocatellia (Acidobacteria group 4); o__Blastocatellales; f__Blastocatellaceae; g__RB41; |
| | OTU_30 | 1 | 0.001 | k__Bacteria; p__Chloroflexi; c__Chloroflexi Subdivision 10; o__Gitt-GS-136; |
| | OTU_60 | 1 | 0.001 | k__Bacteria; p__Actinobacteria; c__Thermoleophilia; o__Gaiellales; f__Gaiellaceae; g__Gaiella; |
| | OTU_19 | 1 | 0.001 | k__Bacteria; p__Chloroflexi; c__Chloroflexi Subdivision 10; o__KD4-96; |
| | OTU_241 | 0.999 | 0.001 | k__Bacteria; p__Acidobacteria; c__Subgroup 6; |
| | OTU_45 | 0.999 | 0.001 | k__Bacteria; p__Actinobacteria; c__Actinobacteria (class); o__Micrococcales; f__Micrococcaceae; |
| | OTU_25 | 0.999 | 0.001 | k__Bacteria; p__Actinobacteria; c__Acidimicrobiia; o__Acidimicrobiales; f__Acidimicrobiaceae; g__CL500-29 marine group; |
| | OTU_75 | 0.999 | 0.001 | k__Bacteria; p__Actinobacteria; c__Thermoleophilia; o__Solirubrobacterales; f__Solirubrobacteraceae; g__Solirubrobacter; |
| | OTU_48300 | 0.999 | 0.001 | k__Bacteria; p__Chloroflexi; c__Chloroflexi Subdivision 10; o__KD4-96; |
| | OTU_64 | 0.999 | 0.001 | k__Bacteria; p__Verrucomicrobia; c__OPB35 soil group; |
| | OTU_240 | 0.998 | 0.001 | k__Bacteria; p__Bacteroidetes; c__Chitinophagia; o__Chitinophagales; f__Chitinophagaceae; |
| | OTU_189 | 0.998 | 0.001 | k__Bacteria; p__Chloroflexi; c__Chloroflexi Subdivision 10; o__KD4-96; |
| | OTU_102 | 0.998 | 0.001 | k__Bacteria; p__Acidobacteria; c__Acidobacteria group 17; |
| | OTU_14100 | 0.998 | 0.001 | k__Bacteria; p__Chloroflexi; c__Chloroflexi Subdivision 10; o__KD4-96; |
| | OTU_224 | 0.998 | 0.001 | k__Bacteria; p__Bacteroidetes; c__Chitinophagia; o__Chitinophagales; f__Chitinophagaceae; g__Ferruginibacter; |
| | OTU_58 | 0.998 | 0.001 | k__Bacteria; p__Acidobacteria; c__Blastocatellia (Acidobacteria group 4); o__Blastocatellales; f__Blastocatellaceae; g__11-24; |
| | OTU_37 | 0.998 | 0.001 | k__Bacteria; p__Acidobacteria; c__Blastocatellia (Acidobacteria group 4); o__Blastocatellales; f__Blastocatellaceae; |
| | OTU_7 | 0.998 | 0.001 | k__Bacteria; p__Acidobacteria; c__Blastocatellia (Acidobacteria group 4); o__Blastocatellales; f__Blastocatellaceae; g__RB41; |
| | OTU_690 | 0.997 | 0.001 | k__Bacteria; p__Verrucomicrobia; c__Spartobacteria; o__Chthoniobacterales; f__DA101 soil group; |

| | | | | |
|---|---|---|---|---|
| | OTU_54 | 0.997 | 0.003 | k__Bacteria; p__Verrucomicrobia; c__Spartobacteria; o__Chthoniobacterales; f__Chthoniobacteraceae; g__Chthoniobacter; |
| | OTU_33 | 0.997 | 0.001 | k__Bacteria; p__Verrucomicrobia; c__Spartobacteria; o__Chthoniobacterales; f__DA101 soil group; |
| | OTU_104 | 0.997 | 0.001 | k__Bacteria; p__Actinobacteria; c__Actinobacteria (class); o__Frankiales; f__Nakamurellaceae; g__Nakamurella; |
| | OTU_68 | 0.997 | 0.001 | k__Bacteria; p__Proteobacteria; c__Betaproteobacteria; o__Burkholderiales; f__Comamonadaceae; |
| | OTU_107 | 0.997 | 0.001 | k__Bacteria; p__Proteobacteria; c__Alphaproteobacteria; o__Rhizobiales; f__Rhizobiales Incertae Sedis; g__Bauldia; |
| | OTU_43 | 0.996 | 0.003 | k__Bacteria; p__Bacteroidetes; c__Chitinophagia; o__Chitinophagales; f__Chitinophagaceae; |
| | OTU_16619 | 0.996 | 0.001 | k__Bacteria; p__Acidobacteria; c__Blastocatellia (Acidobacteria group 4); o__Blastocatellales; f__Blastocatellaceae; g__RB41; |
| | OTU_94 | 0.995 | 0.001 | k__Bacteria; p__Proteobacteria; c__Alphaproteobacteria; o__Rhizobiales; |
| | OTU_36960 | 0.995 | 0.001 | k__Bacteria; p__Acidobacteria; c__Subgroup 6; |
| | OTU_10 | 0.995 | 0.001 | k__Bacteria; p__Gemmatimonadetes; c__Gemmatimonadetes (class); o__Gemmatimonadales; f__Gemmatimonadaceae; |
| | OTU_55 | 0.994 | 0.001 | k__Bacteria; p__Acidobacteria; c__Blastocatellia (Acidobacteria group 4); o__Blastocatellales; f__Blastocatellaceae; g__RB41; |
| | OTU_86 | 0.993 | 0.001 | k__Bacteria; p__Acidobacteria; c__Subgroup 6; |
| | OTU_247 | 0.992 | 0.001 | k__Bacteria; p__Proteobacteria; c__Betaproteobacteria; o__SC-I-84; |
| | OTU_51 | 0.99 | 0.001 | k__Bacteria; p__SPAM; c__; |
| | OTU_21 | 0.989 | 0.001 | k__Bacteria; p__Proteobacteria; c__Betaproteobacteria; o__Nitrosomonadales; f__Nitrosomonadaceae; |
| | OTU_6 | 0.988 | 0.002 | k__Bacteria; p__Actinobacteria; c__Actinobacteria (class); o__Micrococcales; f__Intrasporangiaceae; |
| | OTU_401 | 0.985 | 0.003 | k__Bacteria; p__Acidobacteria; c__Subgroup 6; |
| | OTU_49 | 0.984 | 0.007 | k__Bacteria; p__Acidobacteria; c__Subgroup 6; |
| | OTU_126 | 0.983 | 0.001 | k__Bacteria; p__Gemmatimonadetes; c__Gemmatimonadetes (class); o__Gemmatimonadales; f__Gemmatimonadaceae; |
| | OTU_24 | 0.983 | 0.001 | k__Bacteria; p__Proteobacteria; c__Alphaproteobacteria; o__Rhizobiales; f__MNG7; |

| | | | | |
|---|---|---|---|---|
| | OTU_56 | 0.983 | 0.001 | k__Bacteria; p__Proteobacteria; c__Betaproteobacteria; o__Nitrosomonadales; f__Nitrosomonadaceae; |
| | OTU_72 | 0.979 | 0.005 | k__Bacteria; p__Proteobacteria; c__Alphaproteobacteria; o__Rhizobiales; f__Hyphomicrobiaceae; g__Rhodoplanes; |
| | OTU_99 | 0.979 | 0.001 | k__Bacteria; p__Actinobacteria; c__Actinobacteria (class); o__Pseudonocardiales; f__Pseudonocardiaceae; g__Pseudonocardia; |
| | OTU_21915 | 0.979 | 0.001 | k__Bacteria; p__Acidobacteria; c__Blastocatellia (Acidobacteria group 4); o__Blastocatellales; f__Blastocatellaceae; g__RB41; |
| | OTU_17 | 0.978 | 0.007 | k__Bacteria; p__Acidobacteria; c__Holophagae; o__Acidobacteria group 7a; |
| | OTU_123 | 0.978 | 0.006 | k__Bacteria; p__Proteobacteria; c__Alphaproteobacteria; o__Rhizobiales; f__Xanthobacteraceae; g__Variibacter; |
| | OTU_19327 | 0.974 | 0.003 | k__Bacteria; p__Verrucomicrobia; c__Spartobacteria; o__Chthoniobacterales; f__DA101 soil group; |
| | OTU_95 | 0.966 | 0.001 | k__Bacteria; p__Verrucomicrobia; c__Spartobacteria; o__Chthoniobacterales; f__Chthoniobacteraceae; g__Chthoniobacter; |
| | OTU_20 | 0.966 | 0.001 | k__Bacteria; p__Acidobacteria; c__Blastocatellia (Acidobacteria group 4); o__Blastocatellales; f__Blastocatellaceae; g__RB41; |
| | OTU_31 | 0.948 | 0.005 | k__Bacteria; p__Proteobacteria; c__Betaproteobacteria; o__Nitrosomonadales; f__Nitrosomonadaceae; |
| | OTU_34 | 0.944 | 0.001 | k__Bacteria; p__Proteobacteria; c__Deltaproteobacteria; o__Desulfuromonadales; f__Geobacteraceae; g__Geobacter; |
| | OTU_80 | 0.944 | 0.004 | k__Bacteria; p__Acidobacteria; c__Holophagae; o__Acidobacteria group 7b; |
| | OTU_13683 | 0.944 | 0.005 | k__Bacteria; p__Acidobacteria; c__Blastocatellia (Acidobacteria group 4); o__Blastocatellales; f__Blastocatellaceae; g__RB41; |
| | OTU_50 | 0.931 | 0.027 | k__Bacteria; p__Acidobacteria; c__Blastocatellia (Acidobacteria group 4); o__Blastocatellales; f__Blastocatellaceae; |

1. * OTUs, the relative abundance of which were less than 0.1%, were not listed.
2. ** Taxonomic level: p, phylum; c, class; o, order; f, family; g, genus.

*Table S3: Indicator species determined by the Dufrene-Legendre indicator species analysis method to identify OTUs that are specifically associated with the different pH ranges*
